# A Personalized Metabolic Modelling Approach through Integrated Analysis of RNA-Seq-Based Genomic Variants and Gene Expression Levels in Alzheimer’s Disease

**DOI:** 10.1101/2024.04.24.590807

**Authors:** Dilara Uzuner, Atılay İlgün, Fatma Betül Bozkurt, Tunahan Çakır

## Abstract

**Motivation:** Alzheimer’s disease (AD) is known to cause alterations in brain metabolism. Furthermore, genomic variants in enzyme-coding genes may exacerbate AD-linked metabolic changes. Generating condition-specific metabolic models by mapping gene expression data to genome-scale metabolic models is a routine approach to elucidate disease mechanisms from a metabolic perspective. RNAseq data provides both gene expression and genomic variation information. Integrating variants that perturb enzyme functionality from the same RNAseq data may enhance model accuracy, offering insights into genome-wide AD metabolic pathology.

**Results:** Our study pioneers the extraction of both transcriptomic and genomic data from the same RNA-seq data to reconstruct personalized metabolic models. We mapped genes with significantly higher load of pathogenic variants in AD onto a human genome-scale metabolic network together with the gene expression data. Comparative analysis of the resulting personalized patient metabolic models with the control models showed enhanced accuracy in detecting AD-associated metabolic pathways compared to the case where only expression data was mapped on the metabolic network. Besides, several otherwise would-be missed pathways were annotated in AD by considering the effect of genomic variants.

**Implementation:** The scripts are available at

https://github.com/SysBioGTU/GenomicVariantsMetabolicModels.

## 1. Introduction

Alzheimer’s Disease (AD) is a complex disease without exact treatment. A systems-level perspective is required to discover the molecular mechanisms driving the multifaceted pathogenesis of the disease. Recent research on AD revealed that cellular metabolism is highly affected in the disease. The brain of AD patients shows mitochondrial dysfunction (Bhatia, et al., 2022), impairment of lipid metabolism (Varma, et al., 2021) and reduced overall energy metabolism (Yu, et al., 2022). In the light of these findings, therapeutic strategies for AD have also shifted focus to the metabolism. As an example, the Drug Repurposing for Effective Alzheimer’s Medicines (DREAM) study is a large scale study for drug repurposing on AD and related dementias that target metabolic abnormalities in AD (Desai, et al., 2020).

Genome-scale metabolic models (GEMs) are powerful instruments to study metabolic events in the cell comprehensively. GEMs describe all known metabolic reactions and their gene associations for an organism, enabling the analysis of alterations in metabolic reactions and pathways at different conditions including disease states. Algorithms are available to integrate transcriptomic data with GEMs to generate reduced, condition-specific metabolic models (Gopalakrishnan, et al., 2023). iMAT (Zur, et al., 2010) is one of the most widely used integration algorithms for mammalian cells since it does not require any specific measurement constraints and any definition of biological objective in terms of metabolic reactions, which is the case for the majority of such algorithms. Recently, Baloni et al. (Baloni, et al., 2020) used iMAT to construct AD models and showed that bile acid and cholesterol metabolism reactions were commonly active in different brain-region specific metabolic models. Also, Moolamalla and Vinod (Moolamalla and Vinod, 2020) used the iMAT algorithm to generate metabolic models for different neuropsychiatric diseases like schizophrenia, bipolar disorder, and major depressive disorder.

Whole genome sequencing and whole exome sequencing are popular methods to detect single-nucleotide variants (SNVs) on the genome. RNAseq data can be used to identify pathogenic variants based on the sequencing of exonic regions. This makes RNAseq data unique since both gene expression levels and pathogenic variants can be identified from the same sample (Zhao, 2019). Different studies in the literature have already reported the application of variant discovery from RNAseq data in different diseases (Berger, et al., 2022; Tushir, et al., 2021). However, the use of genomic and transcriptomic information content of RNAseq data together to generate condition-specific genome-scale metabolic models have remained unexplored to date.

In this study, we used the iMAT algorithm to generate personalized genome-scale metabolic models for each individual in three different large AD study cohorts; The Religious Orders Study (ROS) and Rush Memory and Aging Project (MAP) (De Jager, et al., 2018), Mayo Clinic (Allen, et al., 2016) and Mount Sinai Brain Bank (MSBB) (Wang, et al., 2016), which collectively cover a total of 643 AD individuals. For the first time in the literature, we extracted both transcriptomic and genomic information from the same RNA-seq sample to generate personalized metabolic models. We show that the consideration of genes with significantly higher load of pathogenic variants in AD state considerably improves metabolic models by capturing a number of otherwise missed AD-associated metabolic alterations at pathway level.

## 2. Methods

### 2.1. Transcriptome Datasets

Three most commonly used RNAseq-derived AD datasets in the literature were used in this study. ROSMAP, Mayo Clinic and MSBB raw FASTQ files were retrieved from the Synapse Platform (https://www.synapse.org) with accession IDs: syn17024112, syn9738945 and syn7416949, respectively. Clinical metadata of the datasets were also retrieved from the Synapse platform. The ROSMAP dataset was derived from the dorsolateral prefrontal cortex, the Mayo Clinic dataset from the temporal cortex and the MSBB dataset from the parahippocampal gyrus. Samples were categorized as AD and control based on the CERAD score. Individuals with CERAD score 4 (No AD) were considered as the control group, and 1 (Definite AD) and 2 (Probable AD) were considered as AD. Since there was no CERAD score information in the Mayo data, sample classification was done based on Braak Stage (0-III as control, IV-VI as AD). ROSMAP, Mayo Clinic and MSBB RNAseq datasets include 404, 82 and 158 AD and 165, 78 and 26 control samples, respectively.

### 2.2. Gene Expressions from RNAseq data

The FASTQ files were subjected to FastQC tool (version 0.11.9)(Andrews, 2010) for quality control, and low-quality reads, short reads and adapter sequences were trimmed by Trimmomatic (version 0.39) (Bolger, et al., 2014). Trimmed reads were aligned to human reference genome (hg38) by the STAR algorithm (version 2.7.8a) (Dobin, et al., 2013). Then, featureCounts tool (version 2.0.2) (Liao, et al., 2014) was used to obtain raw counts.

Between-sample normalization was applied on the raw counts using Deseq2 (Love, et al., 2014) R package. The log2 values of normalized counts were calculated, and the log-transformed values were adjusted for major covariates (age, gender and post-mortem interval). To this end, the built-in R function *lm* was used to generate linear models and to calculate coefficients of age, gender and post-mortem interval covariates. Subsequently, the effects of the covariates were removed from the normalized and log2-transformed count data.

Principal component analysis (PCA) was performed to detect outlier samples for each dataset. 1 AD (Sample ID: 500_120515) and 1 control (Sample ID: 380_120503) from the ROSMAP dataset and 3 controls from the Mayo Clinic dataset (Sample IDs: 11294, 11396, 11399) were determined to be outliers and were removed from the data.

### 2.3. Variant Identification from RNAseq data

By using the trimmed FASTQ files, the STAR tool was re-run to get splice junction information for each sample. Variant calling was then performed after a second indexing and alignment, using GATK tools (version 4.2.0.0) as described in GATK Best Practices for Variant Calling in RNAseq (Brouard and Bissonnette, 2022). Briefly, AddOrReplaceReadGroups was used to add read group information to BAM files, and it was followed by MarkDuplicates, which identifies duplicate reads resulting from the PCR step. Then, SplitNCigarReads was used to split mapped reads at the intronic regions, and BSQR (Base Quality Score Recalibration) was used to re-score base quality scores. Afterwards, variants were detected using the HaplotypeCaller tool, and a VCF file was generated for each sample. The variants were filtered and annotated by the VariantFiltration tool of GATK and ANNOVAR (Wang, et al., 2010) respectively. In further analyses, biallelic variants with read depth value equal to or greater than five were used.

### 2.4. Gene-level Pathogenicity Score Calculation

The GenePy (Mossotto, et al., 2019) algorithm was employed to transform variant-level pathogenicity scores into gene-level pathogenicity scores. This algorithm operates by aggregating the impact of all variants within a gene in an additive fashion, thereby generating a cumulative pathogenicity score that considers the combined effects of numerous small to moderate effects exerted by each variant. This method closely follows the non-Mendelian inheritance patterns typically observed in complex diseases. GenePy scores of each gene for each individual were computed in the R environment by using the in-house script implemented based on the GenePy score formula and GenePy algorithm.

In this study, Rare Exome Variant Ensembl Learner (REVEL) scores (Ioannidis, et al., 2016) were used as the deleteriousness value. REVEL predicts variant pathogenicity by combining scores from 13 distinct in silico pathogenicity prediction tools. The REVEL score of a variant takes values between 0 and 1, with higher scores reflecting a higher likelihood of the variant being pathogenic. Structural variants (e.g., frameshift indels, stop loss/gain mutations) that lead to protein truncation/elongation are not assigned REVEL scores. Therefore, the maximum deleteriousness value 1 was assigned to all structural variants due to their profoundly damaging impact on protein function. The Genome Aggregation Database (gnomAD) (Karczewski, et al., 2020) exome allele frequencies were used as the allele frequency in the GenePy score calculation. Since the gnomAD exome database encompasses 125,748 individuals, variants lacking frequency data in gnomAD were assigned an allele frequency of 3.98 x 10^−6^, indicating one allele present among 125,748 individuals. The GenePy scores were computed for each sample in each dataset.

Larger genes can accumulate higher GenePy scores as they tend to harbor a greater number of variants, resulting in inflated GenePy scores. To address this, GenePy scores were divided by gene length and multiplied by the median observed gene length in the data for gene length correction (Mossotto, et al., 2019). Finally, the GenePy scores were adjusted to account for gender effect using the *lm* function in R, as mentioned in the previous section. To determine genes with significantly higher pathogenic variants in a personalized manner for each AD individual, the GenePy score of each gene in a given AD sample was combined with the scores of that gene in all the control samples in that dataset, and the values were ranked. If the score is higher than 95% of the scores from the control individuals, the gene was marked as a gene with higher load of pathogenic variants in AD.

### 2.5. Personalized Metabolic Model Reconstruction

To reconstruct personalized genome-scale metabolic models for all controls and AD cases, genes in the covariate-adjusted expression data were mapped individually to the human genome-scale metabolic model (Human-GEM) using the integrative metabolic analysis tool (iMAT) (Zur, et al., 2010). The iMAT algorithm was run with the parameters described in our previous study (Lüleci, et al., 2023). Due to the covariate adjustment of the expression data, there are negative values in the data. iMAT algorithm available through the COBRA Toolbox (Heirendt, et al., 2019) ignores genes with negative expression values. Hence, the absolute value of the smallest negative expression value in each dataset was added to the whole data. In this way, all expression values were converted to positive values. Since the parameters used by the iMAT algorithm to determine active and inactive reactions are based on percentiles, this process does not affect the results. Human-GEM (version 1.12.0) (Wang, et al., 2022) was retrieved from the GitHub repository (https://github.com/SysBioChalmers/Human-GEM). This comprehensive metabolic network includes 13,070 reactions associated with 3,067 genes, 8,369 metabolites and 143 pathways. 3,055 out of 3,067 genes in the Human-GEM are included in the transcriptome datasets. Simulations were performed using MATLAB R2020a with the Gurobi solver.

To incorporate the effects of the variants that disrupt protein function into personalized metabolic models (Figure 1A), the following steps were followed: (i) It was hypothesized that genes with significantly higher load of pathogenic variants in AD, as identified by the GenePy algorithm, would not be able to form functional proteins due to mutations on them, even if their mRNA expression levels were not low. Based on this assumption, the expression levels of pathogenic variant genes were set to zero in the covariate adjusted expression data of that AD sample. This was repeated for all AD samples. (ii) Then, sample-based iMAT models were generated by forcing the removal of the reactions controlled by the combined list of the lowly expressed genes and the genes with pathogenic variants from the metabolic model, subject to mass-balance constraints around the intracellular metabolites. iMAT restores the reactions back if they violate mass-balance constraints or if the removal decreases the consistency of reaction rates with the transcriptome data. The resulting metabolic models were referred as AD^var^ models in this study.

**Figure 1:**
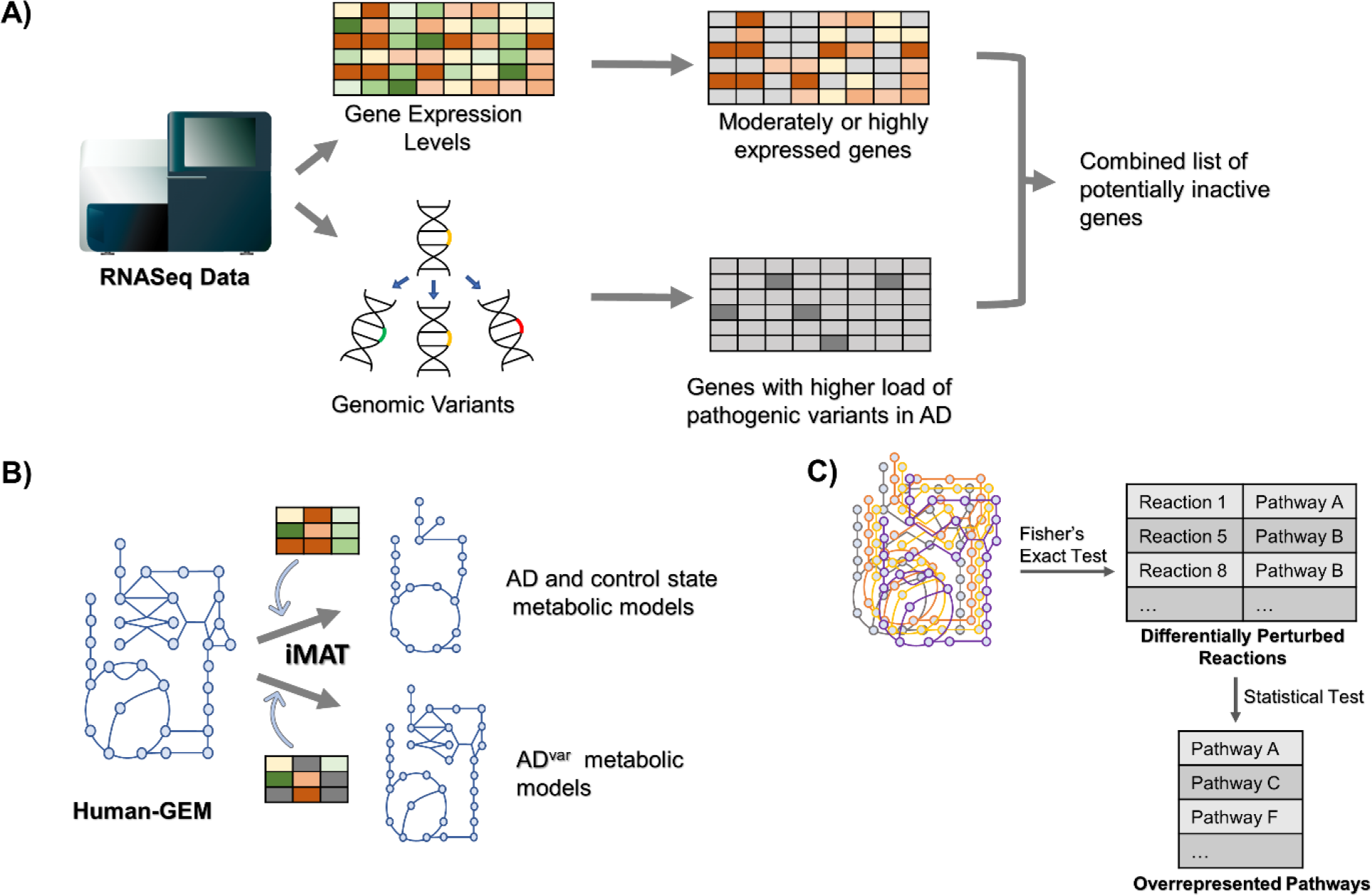
Summary of iMAT-based personalized metabolic model generation and incorporation of the impact of pathogenic variants to models. **A)** RNAseq data are obtained in Fastq format and processed with two different pipelines to find gene expression values and genomic variants. The genes with higher load of pathogenic variants and the genes with high or moderate expression levels were selected as inactive genes for each AD sample. **B)** Gene expression values are mapped to the genome-scale metabolic model and AD, control and AD^var^ models are created with the iMAT algorithm. To create AD^var^ models, unlike the classical method, the genes carrying higher load of pathogenicity in AD were additionally used. **C)** Differentially perturbed reactions between AD and control are identified by Fisher’s Exact Test after representing each personalized metabolic model as a binary vector, where “1” means the reaction is active in that model. Then, pathways overrepresented with the differentially perturbed reactions were predicted.

### 2.6. Statistical Analyses

The generated iMAT models were converted to binary format to apply the statistical tests. This conversion was based on the reactions in Human-GEM. Reactions active in the iMAT models were represented by 1 and inactive reactions were represented by 0 for each sample. Then, using these binary matrices, a contingency table was created for each reaction between AD and control conditions, and Fisher’s Exact Test was performed to identify significantly altered reactions in each dataset. P-value cut-off of 0.05 was used for AD-Control models while the cut-off was 0.01 for AD^var^-Control models.

The reaction-subsystem assignments in Human-GEM were used to find significantly affected pathways. Fisher Exact Test based p-values of each reaction were converted to z-scores using the inverse cumulative density function. Each pathway z-score was then calculated by averaging the z-scores of the reactions associated with that pathway and corrected with 1000 random permutations as described elsewhere (Çakır, 2015; Patil and Nielsen, 2005). Corrected pathway z-scores were converted to pathway p-values using the cumulative density function. Significantly affected pathways were identified using the p-value < 0.05 cut-off. This approach enabled consideration of all pathway reactions in the calculation compared to a hypergeometric test where only reactions with significant p-values are considered.

### 2.7. Identification of AD-Related Genes Associated with the Affected Metabolic Reactions

Using the Gene-Protein-Reaction (GPR) rules of Human-GEM, the genes controlling significantly perturbed reactions were extracted. Experimentally confirmed and curated AD-related gene lists were collected from another study (Ceylan, et al., 2024) and DisGeNET (Piñero, et al., 2021). There are 166 AD-related metabolic genes in the final list. The hypergeometric test was used to test whether the number of AD-related genes was significantly overrepresented in the identified lists of significantly perturbed reactions from AD or AD^var^ models.

## 3. Results and Discussion

### 3.1. iMAT Models of AD, AD^var^ and Control conditions

Each sample in the ROSMAP, Mayo Clinic and MSBB post-mortem brain RNAseq datasets were mapped to Human-GEM using the iMAT algorithm to reconstruct personalized genome-scale metabolic models for both AD and control samples. This led to 908 models in total spanning the three datasets.

The iMAT algorithm generates metabolic models by inactivating reactions controlled by enzymes encoded by genes with low expression, while keeping reactions controlled by enzymes encoded by genes with high expression in the model, subject to mass-balance constraints around intracellular metabolites (Zur, et al., 2010). It also ensures maximum consistency between gene expression levels and reaction rates. On the other hand, some genomic variants are pathogenic, i.e. they lead to dysfunctional enzymes, although they do not affect expression at the mRNA level (Chen, et al., 2004; Lockett, et al., 2004). In this study, first, genomic variants in the form of single-nucleotide variants and insertions/deletions were predicted from the RNAseq datasets, and, then, their in silico pathogenicity scores were used to calculate gene-level scores. The correlation between expression levels and pathogenicity scores of metabolic genes was calculated for each sample. The correlation coefficients ranged between -0.03-0.23 for all the samples from three different cohorts (Supplementary Figure 1). Therefore, the pathogenicity scores of each sample do not depend on gene expression levels and provide a novel knowledge. To integrate this knowledge into metabolic models, genes with significantly higher pathogenicity in AD samples were marked as inactive in generating iMAT-derived metabolic models, termed AD^var^ models (Figure 1A-B).

The number of metabolic genes carrying pathogenic variants in AD samples was about 50-150 for the studied datasets (Figure 2A). This means that the genes additionally included in the list of genes to be removed from the metabolic models to create AD^var^ models is about ∼1.5%-5% of the total number of genes in Human-GEM, which is not a high fraction. In other words, our approach of incorporating genes with higher load of pathogenic variants in AD introduces minimal intervention to the original iMAT approach. We also checked the expression levels of these genes to see how many of them are expressed higher than the low-expression cut-off used by iMAT. On average, we found 3 pathogenic variant carrying genes in ROSMAP, 30 in Mayo and 4 in MSBB to have expression levels higher than 25% of all the genes in transcriptome data (Figure 2B). Thus, the number of metabolic genes manipulated in the transcriptome data is considerably less than the number of genes in the model.

**Figure 2:**
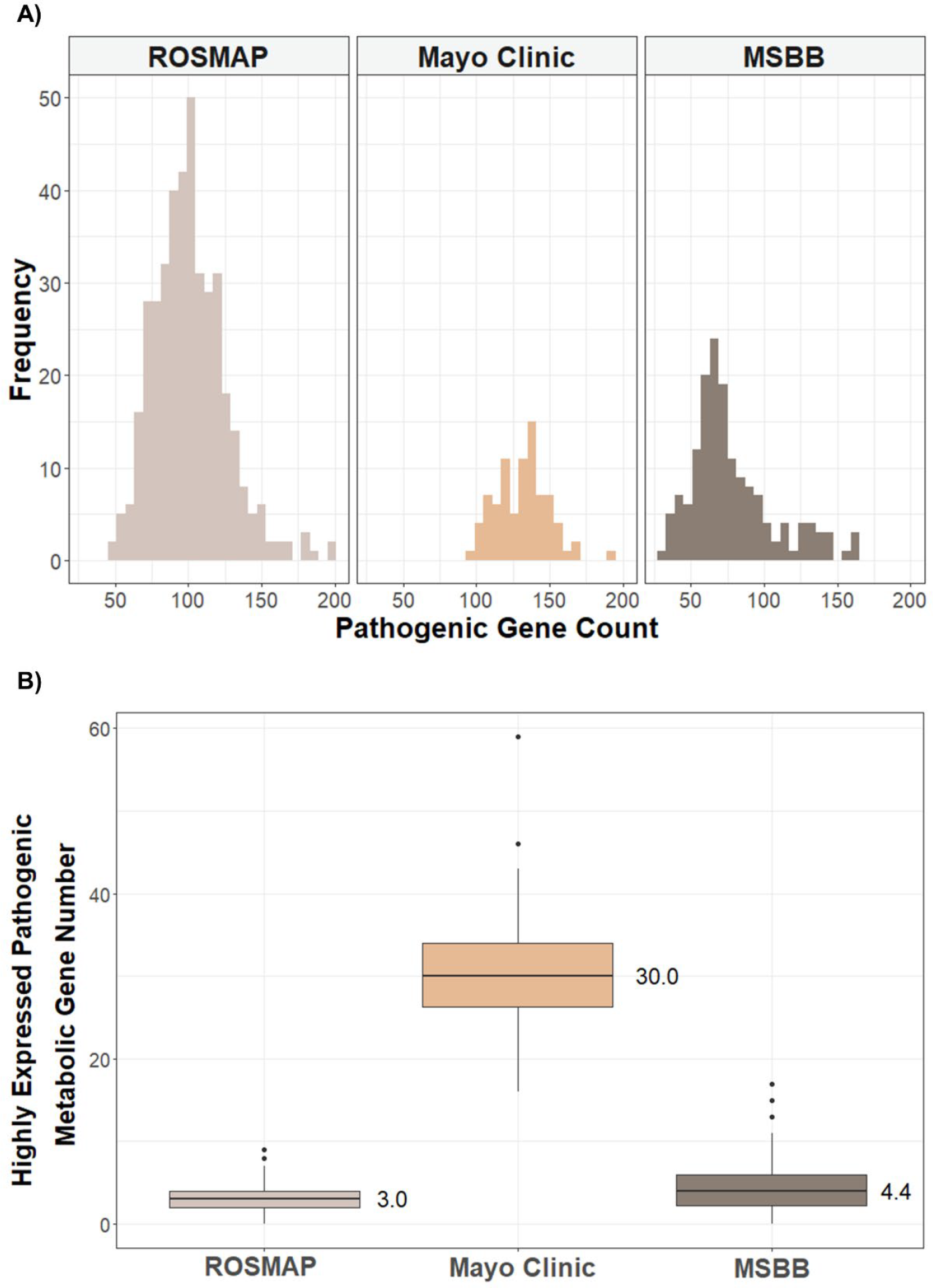
Number of metabolic genes carrying significantly higher load of pathogenic variants in AD for each dataset. **A)** Histogram of the number of metabolic genes with higher load of pathogenic variants in each AD sample for each dataset. Frequency axis shows the number of samples having a specific number of pathogenic genes as indicated in the x-axis. **B)** Boxplot of the number of metabolic genes carrying pathogenic variants and having expression levels higher than 25% of the genes (the iMAT low-expression cut-off) in the whole transcriptome.

Subsequently, the iMAT models of each dataset were compared to investigate whether the numbers of active reactions between the control, AD^var^ and AD models are different. As shown in Figure 3A, all models consisted of ∼7000-8000 reactions while Human-GEM includes 13070 reactions. Accordingly, it was observed that the distribution of the number of active reactions was more similar in the AD and control models compared to the AD^var^ models. In the AD^var^ models, although there was no dramatic difference in terms of the number of reactions in the created personalized models, the distribution became more spread out. For each sample, we also looked at the difference between the number of reactions between the AD and AD^var^ models of that sample (Figure 3B) and found that there was a difference of about 50 reactions in ROSMAP and MSBB and about 200 reactions for Mayo Clinic. Differing from the other datasets, the AD^var^ metabolic models of the Mayo Clinic dataset had fewer reactions than the AD models. To better understand this difference, Jaccard similarity indices were calculated between the binary matrices representing AD and AD^var^ models for each dataset, and the similarity of the binary models was around 85% for all the datasets. These results indicate that although there are differences between the models at the individual level, in general the AD and AD^var^ models are similar in terms of the majority of active reactions.

**Figure 3:**
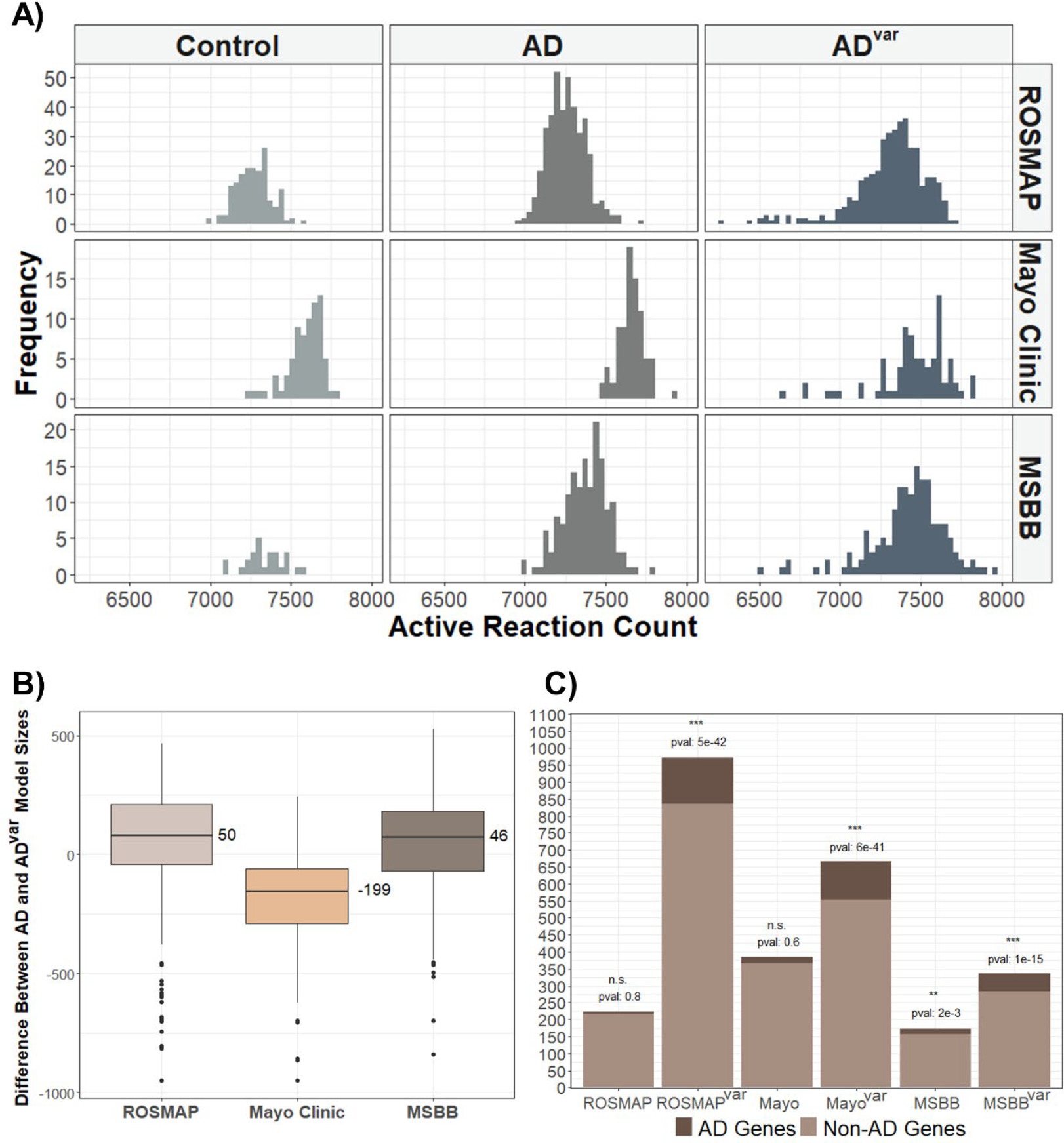
Condition Specific Model Generation By iMAT. **A)** Histogram of the number of active reactions in each personalized model for each dataset. Frequency shows the number of instances with the count. **B)** Boxplot of the difference in reaction numbers between AD and AD^var^ iMAT models for each sample. **C)** Bar plot showing the total number of genes associated with the lists of significantly altered reactions, and the proportion of AD-related genes in these lists for each dataset and for each of AD and AD^var^ models. pval shows the hypergeometric p-value obtained from the overrepresentation analysis of AD-related genes in each list. n.s: not significant, ** pval<0.01, ***pval<0.001.

After the significantly altered reactions were identified using Fisher’s Exact test, gene compositions controlling these reactions were analyzed to determine the number of genes associated with AD for each reaction list (Figure 4). Accordingly, while the significant reaction lists were significantly enriched with the AD-related genes was in all AD^var^ models, a significant result was obtained only in MSBB among the models constructed with only gene-expression data, and with a much higher p-value compared to the AD^var^ counterpart. This shows that the integration of pathogenic-variant data into the metabolic models contributes to a better representation of AD status at the gene level.

**Figure 4:**
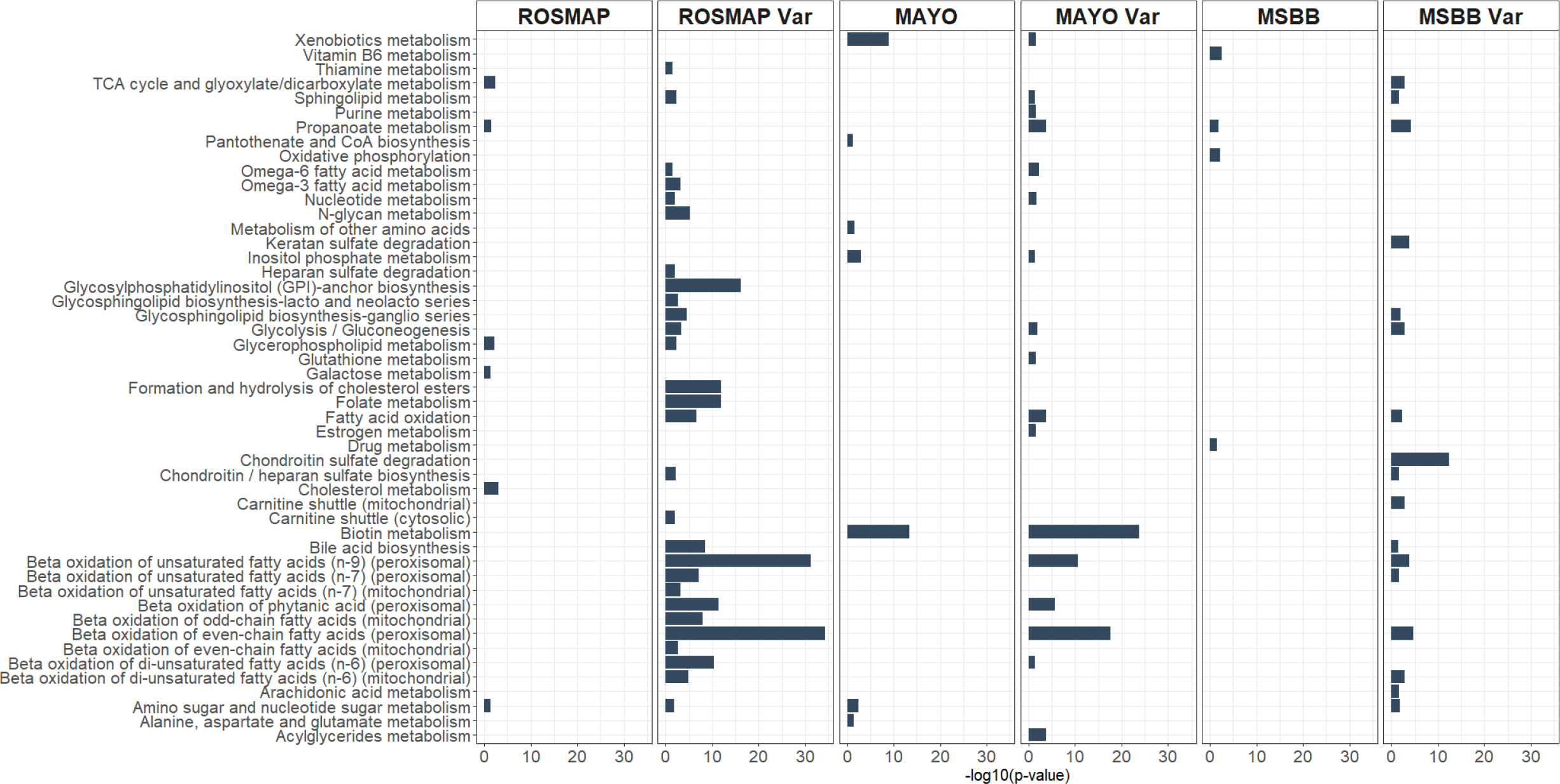
Differentially perturbed pathways between AD-Control models and AD^var^-Control (Var) models. Pathway enrichment analysis was applied on the differentially altered reactions. x-axis shows -log10(p-value) of the pathways. Only significant pathways are shown in the figure for each dataset.

### 3.2. Differentially Perturbed Pathways

After the personalized models were generated, Fisher’s Exact Test was applied to identify differentially affected reactions between AD-Control comparison and AD^var^-Control comparison (Figure 1C, Supplementary File 2). Although the model sizes and pathogenic gene counts were not much different in the AD and AD^var^ models, the number of differentially perturbed reactions between AD-Control and AD^var^-Control were considerably different (Supplementary Figure 2). For all the cohorts, adding the effect of pathogenic variants to the models increased the number of differentially altered reactions (Table 1). Using reaction-pathway associations of Human-GEM, we also identified the number of unique pathways associated with the significantly affected reactions for each dataset (Table 1, Supplementary Figure 3).

**Table 1:** The number of significantly affected reactions and associated pathways.

| **Datasets and Compared Conditions** | **# of significantly affected reactions** | **# of associated pathways** | **# of overrepresented pathways** |
| --- | --- | --- | --- |
| **ROSMAP AD-Control** | 290 | 55 | 6 |
| **ROSMAP AD<sup>var</sup>-Control** | 3089 | 108 | 28 |
| **Mayo Clinic AD-Control** | 674 | 64 | 7 |
| **Mayo Clinic AD<sup>var</sup>-Control** | 1343 | 98 | 17 |
| **MSBB AD-Control** | 220 | 40 | 4 |
| **MSBB AD<sup>var</sup>-Control** | 793 | 72 | 17 |

We applied a statistical analysis to identify pathways significantly overrepresented among the affected reactions (Figure 4, Table 1). In general, only few pathways were identified to be significantly perturbed based on the AD models, regardless of the dataset used. We identified a number of pathways whose perturbation was only captured with the AD^var^ models. Below, we provide a detailed analysis of the pathways identified to be significantly perturbed only in AD^var^ models.

The sphingolipid metabolism was significantly perturbed only in the AD^var^ models in all the three cohorts. Sphingolipids are a member of lipid family, and they are crucial elements of membrane biology and have roles in the regulation of cell function (Hannun and Obeid, 2018). Sphingolipid metabolism was previously associated with dementia and the AD risk factor gene APOE genotype (Alaamery, et al., 2021). Also, sphingomyelin metabolism was recently proposed as a therapeutic target for AD based on multi-omics profiling of the AD brain (Baloni, et al., 2022). Therefore, identifying sphingolipid metabolism only in AD^var^ models shows the increased accuracy in capturing AD-related processes in the metabolic models.

Glycolysis/glucogenesis is another pathway that was significantly affected only in AD^var^ models in the three cohorts. Glucose metabolism is important in energy production, which is vital for all cells, and it was reported that glucose metabolism and glycolysis defects contribute to AD pathogenesis (Bergau, et al., 2019; Pan, et al., 2022; Theurey, et al., 2019). In the brain, astrocyte cells prefer aerobic glycolysis to produce lactate. Lactate produced from glycolysis or glycogenolysis in astrocytes is transferred to neurons to be used in oxidative phosphorylation. However, it was observed that the lactate production capacity of astrocytes decreased in AD. (Wang, et al., 2022). Consequently, energy deficiency in neurons and glia play an essential role in AD.

Fatty acid oxidation is another pathway that was significantly affected only in AD^var^ models in the three cohorts. Brain is the organ where fatty acids are the most abundant in our body. It was shown that the levels of all subclasses of fatty acids are different in the cerebrospinal fluid of AD patients compared to controls (Yin, 2023). Qi et al. reported that APOE4 genotype caused a shift in both glucose and fatty acid oxidation in astrocytes (Qi, et al., 2021). Overall, these results indicate that integrating pathogenic variants into the models allows capturing of some important AD-related pathways in different cohorts.

In models generated with the samples obtained from different brain regions, there were also differences among significantly perturbed pathways that were captured only in the variant-aware models. Bile acid, chondroitin/heparan sulfate and glycosphingolipid biosynthesis were differentially perturbed only in the variant models (MSBB and ROSMAP cohorts). Among these three pathways, bile acid metabolism is highly integrated with cholesterol metabolism and has been associated with AD. Recently, Baloni et al. used metabolic modelling approach and reported that bile acid metabolism was altered in AD brains (Baloni, et al., 2020). Similarly, Varma et al. showed that altered brain bile acid metabolism was associated with AD (Varma, et al., 2021). Heparan sulphate is another important metabolic pathway affecting amyloid-beta clearance, and high levels of heparan sulphate proteoglycans were found in post-mortem AD brains (Liu, et al., 2016; Ozsan McMillan, et al., 2023). Glycosphingolipids are a heterogeneous group of membrane lipids. They comprise a high amount of the brain lipid composition. Tang et al. (Tang, et al., 2023) analysed the expression levels of glycosylation related genes in the AD brains and found alterations in the levels of the genes that play a role in glycosphingolipid biosynthesis in human.

Similarly, omega-6 fatty acid, fatty acid beta-oxidation and nucleotide metabolisms were differentially perturbed only in the variant models of Mayo Clinic and ROSMAP. Lipid metabolism including fatty acids is one of the most studied metabolic pathways in AD (Chew, et al., 2020; Kao, et al., 2020). Lipid metabolism is known to be impaired in the early stages of AD (Yin, 2023). On the other hand, since nucleotide metabolism is closely related to cell proliferation pathways, it is interesting that it was also captured in our results although it is a pathway generally studied in cancer (Wu, et al., 2022). As a result, different datasets have the capacity to capture different pathways that have been associated with AD or that may be candidates for AD. These results are also important for understanding how different brain regions are affected in AD.

Additionally, propanoate, inositol phosphate, glycerophospholipid and biotin metabolic pathways were identified to be perturbed in both variant and non-variant models of the same dataset. Propanoate metabolite is known to be neuroprotective and associated with brain-gut axis (Bayraktar, et al., 2021). A study showed that changes in the levels of inositol phosphate metabolic enzymes were correlated with amyloid beta peptide formation and tau hyperphosphorylation in AD (Stygelbout, et al., 2014). A recent study using Drosophila model of tauopathy suggested biotin metabolism as druggable target for neuroprotection (Lohr, et al., 2020). Glycerophospholipids are complex species of fatty acids, and they are the most abundant class of lipids in the human brain (Garcia Corrales, et al., 2021). These results also indicated that adding the variant effect to the models does not cause the loss of previously annotated metabolic processes.

## 4. Conclusion

Genome-scale metabolic models in complex diseases such as Alzheimer’s disease, in which many different metabolisms in the cell are affected, provide a great advantage for systemic investigation of the etiology of the disease. For this purpose, the generation of disease models by integrating gene expression data into GEMs has become widespread in the literature in the last 20 years, and it is a frequently used method today. In these models, information from mRNA level is incorporated into the model as the abundance of enzyme-coding proteins, and optimization problems based on mass-balance around intracellular metabolites are solved by aiming to keep the reactions catalyzed by enzymes with high abundance in the model and excluding the reactions catalyzed by enzymes with low abundance from the model. Although these models are very convenient, possible loss of function in the mRNA-to-protein transition is neglected. In this study, for the first time in the literature, we constructed personalized metabolic models by determining both gene expression levels and pathogenic variants from the same AD and control RNAseq samples from different brain regions. When we compared these models with the models constructed with only gene-expression data, we showed that pathways such as fatty acid oxidation, bile acid metabolism, sphingolipid metabolism and glycolysis, which are known to play crucial roles in AD metabolism, were found to be significantly altered only in the variant integrated models. In addition, gene-level analysis showed that the list of significantly perturbed reactions were significantly enriched with AD genes for all the three datasets in the AD^var^ models, and with a much higher significance level compared to the models constructed with only gene expression data.

Although the results of this study have brought a new perspective to the use of metabolic modelling and transcriptome data in AD studies, it has some limitations. Within the framework of this study, only the effect of genomic variants that are predicted to affect protein expression was considered. However, some variants in intronic regions called eQTLs are known to regulate the expression of target genes. Transcriptome-wide association studies (TWAS) aim to reveal such variant-expression-trait relationships (Li and Ritchie, 2021). In future studies, metabolic models can be generated by also adding the information of metabolic genes whose expressions are regulated by variants based on TWAS studies. In addition, post-translational modifications which could have an important role in enzyme activity were ignored in this study. For more comprehensive results, integrating the effect of post-translational modifications in the cell into the genome-scale metabolic models is another important future study.

## Acknowledgements

This study was financially supported by a grant from The Scientific and Technological Research Council of Türkiye (TUBITAK) (Project Code: 120S824). Also, Dilara Uzuner is funded by TUBITAK 2211-A PhD Student Scholarship and Council of Higher Education (CoHE) 100/2000 PhD Scholarship Programmes.

## References

Alaamery, M., et al. Role of sphingolipid metabolism in neurodegeneration. Journal of Neurochemistry 2021;158(1):25–35.

Allen, M., et al. Human whole genome genotype and transcriptome data for Alzheimer’s and other neurodegenerative diseases. Scientific Data 2016;3.

Andrews, S. FastQC: a quality control tool for high throughput sequence data. In.: Babraham Bioinformatics, Babraham Institute, Cambridge, United Kingdom; 2010.

Baloni, P., et al. Multi-Omic analyses characterize the ceramide/sphingomyelin pathway as a therapeutic target in Alzheimer’s disease. Communications Biology 2022;5(1):1074.

Baloni, P., et al. Metabolic Network Analysis Reveals Altered Bile Acid Synthesis and Metabolism in Alzheimer’s Disease. Cell Rep Med 2020;1(8):100138.

Bayraktar, A., et al. Revealing the Molecular Mechanisms of Alzheimer’s Disease Based on Network Analysis. Int J Mol Sci 2021;22(21).

Bergau, N., et al. Reduction of glycolysis intermediate concentrations in the cerebrospinal fluid of Alzheimer’s disease patients. Frontiers in neuroscience 2019;13:871–871.

Berger, K., et al. Targeted RNAseq Improves Clinical Diagnosis of Very Early-Onset Pediatric Immune Dysregulation. In, Journal of Personalized Medicine. 2022.

Bhatia, S., et al. Mitochondrial Dysfunction in Alzheimer’s Disease: Opportunities for Drug Development. Curr Neuropharmacol 2022;20(4):675–692.

Bolger, A.M., Lohse, M. and Usadel, B. Trimmomatic: a flexible trimmer for Illumina sequence data. Bioinformatics 2014;30(15):2114–2120.

Brouard, J.-S. and Bissonnette, N. Variant Calling from RNA-seq Data Using the GATK Joint Genotyping Workflow. In: Ng, C.and Piscuoglio, S., editors, Variant Calling: Methods and Protocols. New York, NY: Springer US; 2022. p. 205–233.

Ceylan, B., Düz, E. and Çakir, T. Personalized Protein-Protein Interaction Networks Towards Unraveling the Molecular Mechanisms of Alzheimer’s Disease. Molecular Neurobiology 2024;61(4):2120–2135.

Chen, J., et al. Functional analysis of genetic variation in catechol-O-methyltransferase (COMT): effects on mRNA, protein, and enzyme activity in postmortem human brain. The American Journal of Human Genetics 2004;75(5):807–821.

Chew, H., Solomon, V.A. and Fonteh, A.N. Involvement of Lipids in Alzheimer’s Disease Pathology and Potential Therapies. Front Physiol 2020;11:598.

Çakir, T. Reporter pathway analysis from transcriptome data: metabolite-centric versus reaction-centric approach. Scientific Reports 2015;5(1):14563.

De Jager, P.L., et al. A multi-omic atlas of the human frontal cortex for aging and Alzheimer’s disease research. Scientific Data 2018;5(1):180142–180142.

Desai, R.J., et al. Targeting abnormal metabolism in Alzheimer’s disease: The Drug Repurposing for Effective Alzheimer’s Medicines (DREAM) study. Alzheimer’s & Dementia: Translational Research & Clinical Interventions 2020;6(1):e12095.

Dobin, A., et al. STAR: ultrafast universal RNA-seq aligner. Bioinformatics 2013;29(1):15–21.

Garcia Corrales, A.V., et al. Fatty Acid Synthesis in Glial Cells of the CNS. International journal of molecular sciences 2021;22(15):8159.

Gopalakrishnan, S., et al. Guidelines for extracting biologically relevant context-specific metabolic models using gene expression data. Metabolic Engineering 2023;75:181–191.

Hannun, Y.A. and Obeid, L.M. Sphingolipids and their metabolism in physiology and disease. Nature Reviews Molecular Cell Biology 2018;19(3):175–191.

Heirendt, L., et al. Creation and analysis of biochemical constraint-based models using the COBRA Toolbox v.3.0. Nat Protoc 2019;14(3):639–702.

Ioannidis, N.M., et al. REVEL: an ensemble method for predicting the pathogenicity of rare missense variants. The American Journal of Human Genetics 2016;99(4):877–885.

Kao, Y.-C., et al. Lipids and Alzheimer’s Disease. In, International Journal of Molecular Sciences. 2020.

Karczewski, K.J., et al. The mutational constraint spectrum quantified from variation in 141,456 humans. Nature 2020;581(7809):434–443.

Li, B. and Ritchie, M.D. From GWAS to Gene: Transcriptome-Wide Association Studies and Other Methods to Functionally Understand GWAS Discoveries. Frontiers in Genetics 2021;12.

Liao, Y., Smyth, G.K. and Shi, W. featureCounts: an efficient general purpose program for assigning sequence reads to genomic features. Bioinformatics 2014;30(7):923–930.

Liu, C.-C., et al. Neuronal heparan sulfates promote amyloid pathology by modulating brain amyloid-β clearance and aggregation in Alzheimer’s disease. Science Translational Medicine 2016;8(332):332ra344–332ra344.

Lockett, K.L., et al. The ADPRT V762A Genetic Variant Contributes to Prostate Cancer Susceptibility and Deficient Enzyme Function. Cancer research 2004;64(17):6344–6348.

Lohr, K.M., et al. Biotin rescues mitochondrial dysfunction and neurotoxicity in a tauopathy model. Proceedings of the National Academy of Sciences 2020;117(52):33608–33618.

Love, M., Anders, S. and Huber, W. Differential analysis of count data–the DESeq2 package. Genome Biol 2014;15(550):10–1186.

Lüleci, H.B., et al. Computational Approaches to Assess Abnormal Metabolism in Alzheimer’s Disease Using Transcriptomics. In: Chun, J., editor, Alzheimer’s Disease: Methods and Protocols. New York, NY: Springer US; 2023. p. 173–189.

Moolamalla, S.T.R. and Vinod, P.K. Genome-scale metabolic modelling predicts biomarkers and therapeutic targets for neuropsychiatric disorders. Computers in Biology and Medicine 2020;125:103994.

Mossotto, E., et al. GenePy - a score for estimating gene pathogenicity in individuals using next-generation sequencing data. BMC Bioinformatics 2019;20(1):254.

Mossotto, E., et al. GenePy - a score for estimating gene pathogenicity in individuals using next-generation sequencing data. BMC Bioinformatics 2019;20(1):254–254.

Ozsan McMillan, I., Li, J.-P. and Wang, L. Heparan sulfate proteoglycan in Alzheimer’s disease: aberrant expression and functions in molecular pathways related to amyloid-β metabolism. American Journal of Physiology-Cell Physiology 2023;324(4):C893–C909.

Pan, R.-Y., et al. Positive feedback regulation of microglial glucose metabolism by histone H4 lysine 12 lactylation in Alzheimer’s disease. Cell Metabolism 2022;34(4):634-648.e636.

Patil, K.R. and Nielsen, J. Uncovering transcriptional regulation of metabolism by using metabolic network topology. Proceedings of the National Academy of Sciences of the United States of America 2005;102(8):2685.

Piñero, J., et al. The DisGeNET cytoscape app: Exploring and visualizing disease genomics data. Computational and Structural Biotechnology Journal 2021;19:2960–2967.

Qi, G., et al. ApoE4 Impairs Neuron-Astrocyte Coupling of Fatty Acid Metabolism. Cell Reports 2021;34(1).

Stygelbout, V., et al. Inositol trisphosphate 3-kinase B is increased in human Alzheimer brain and exacerbates mouse Alzheimer pathology. Brain 2014;137(2):537–552.

Tang, X., et al. Transcriptomic and glycomic analyses highlight pathway-specific glycosylation alterations unique to Alzheimer’s disease. Scientific Reports 2023;13(1):7816.

Theurey, P., et al. Systems biology identifies preserved integrity but impaired metabolism of mitochondria due to a glycolytic defect in Alzheimer’s disease neurons. Aging Cell 2019;18(3):e12924–e12924.

Tushir, S., et al. Proteo-Genomic Analysis of SARS-CoV-2: A Clinical Landscape of Single-Nucleotide Polymorphisms, COVID-19 Proteome, and Host Responses. Journal of Proteome Research 2021;20(3):1591–1601.

Varma, V.R., et al. Abnormal brain cholesterol homeostasis in Alzheimer’s disease—a targeted metabolomic and transcriptomic study. npj Aging and Mechanisms of Disease 2021;7(1):11.

Varma, V.R., et al. Bile acid synthesis, modulation, and dementia: A metabolomic, transcriptomic, and pharmacoepidemiologic study. PLoS Med 2021;18(5):e1003615.

Wang, H., et al. SysBioChalmers/Human-GEM: Human 1.12.0. In.: Zenodo; 2022.

Wang, K., Li, M. and Hakonarson, H. ANNOVAR: functional annotation of genetic variants from high-throughput sequencing data. Nucleic Acids Research 2010;38(16):e164–e164.

Wang, M., et al. Integrative network analysis of nineteen brain regions identifies molecular signatures and networks underlying selective regional vulnerability to Alzheimer’s disease. Genome Med 2016;8(1):104.

Wang, Q., et al. Glucose Metabolism, Neural Cell Senescence and Alzheimer’s Disease. In, International Journal of Molecular Sciences. 2022.

Wu, H.-l., et al. Targeting nucleotide metabolism: a promising approach to enhance cancer immunotherapy. Journal of Hematology & Oncology 2022;15(1):45.

Yin, F. Lipid metabolism and Alzheimer’s disease: clinical evidence, mechanistic link and therapeutic promise. The FEBS Journal 2023;290(6):1420–1453.

Yu, L., et al. Aberrant Energy Metabolism in Alzheimer’s Disease. J Transl Int Med 2022;10(3):197–206.

Zhao, S. Alternative splicing, RNA-seq and drug discovery. Drug Discovery Today 2019;24(6):1258–1267.

Zur, H., Ruppin, E. and Shlomi, T. iMAT: an integrative metabolic analysis tool. Bioinformatics 2010;26(24):3140–3142.

