## Supplementary Figures for "A Personalized Metabolic Modelling Approach through Integrated Analysis of RNA-Seq-Based Genomic Variants and Gene Expression Levels in Alzheimer’s Disease"

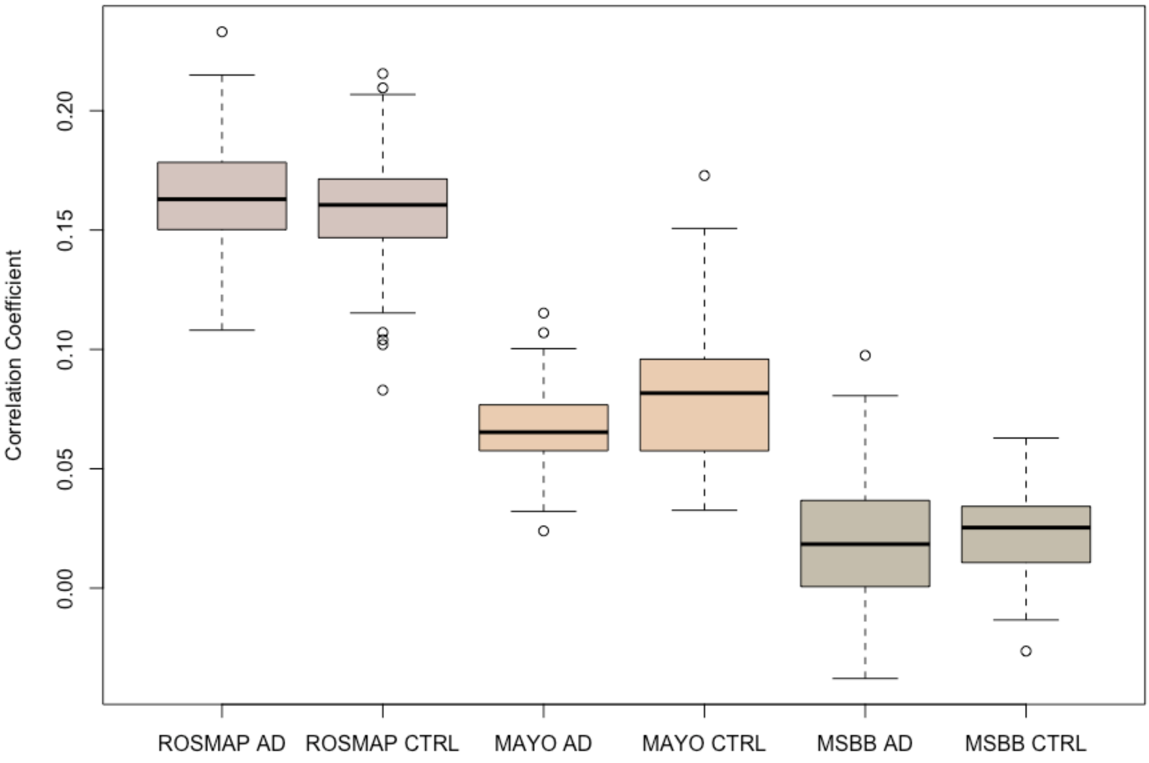


**Supplementary Figure 1: Correlation between expression levels of metabolic genes and gene pathogenicity scores.** The correlation between gene expression levels and gene pathogenicity scores for metabolic genes in ROSMAP, Mayo Clinic and MSBB datasets were calculated separately for each AD and control sample using Pearson correlation method with "cor.test" function in R. Significant correlation ($p<0.05$) was observed in most of the samples except for 1/403, (ROSMAP AD), 4/82 and 3/75 (Mayo AD and Control), 127/158 and 23/26 (MSBB AD and Control) samples have insignificant correlation ($p>0.05$).

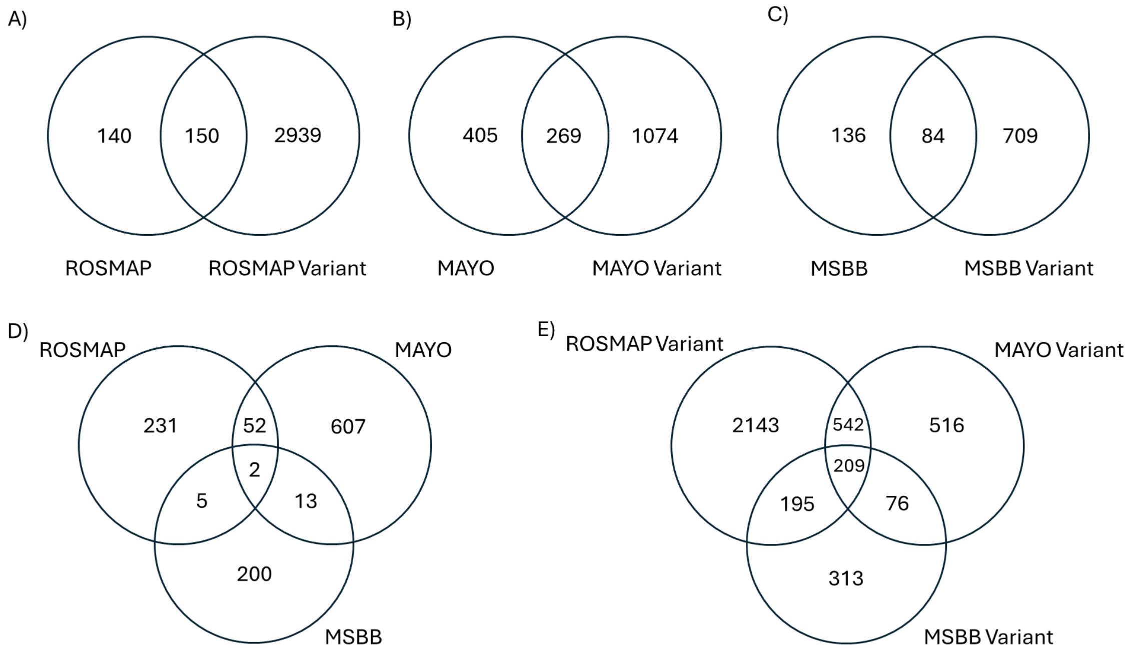


**Supplementary Figure 2**: Venn diagrams of the differentially altered reactions of iMAT models. **A)** ROSMAP AD vs Control and $AD^{var}$ vs Control, **B)** Mayo Clinic AD vs Control and $AD^{var}$ vs Control, **C)** MSBB AD vs Control and $AD^{var}$ vs Control, **D)** ROSMAP, Mayo Clinic and MSBB AD vs Control, **E)** ROSMAP, Mayo Clinic and MSBB $AD^{var}$ vs Control

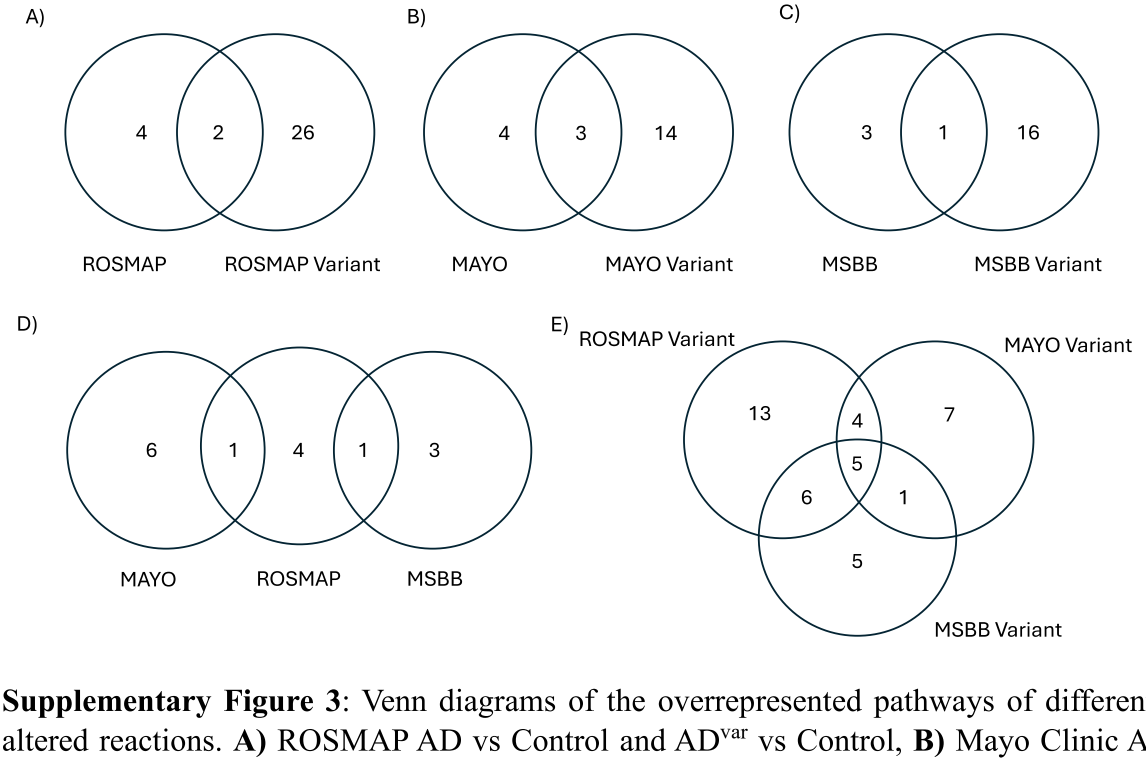


**Supplementary Figure 3**: Venn diagrams of the overrepresented pathways of differentially altered reactions. **A)** ROSMAP AD vs Control and $AD^{var}$ vs Control, **B)** Mayo Clinic AD vs Control and $AD^{var}$ vs Control, **C)** MSBB AD vs Control and $AD^{var}$ vs Control, **D)** ROSMAP, Mayo Clinic and MSBB AD vs Control, **E)** ROSMAP, Mayo Clinic and MSBB $AD^{var}$ vs Control
